# Snake Fungal Disease Caused by the Fungal Pathogen *Ophidiomyces ophidiicola* in Texas

**DOI:** 10.1101/2022.04.14.488407

**Authors:** Alan Lizarraga, Lezley Hart, Michele Nolen, Lance Williams, Joseph Glavy

**Affiliations:** Biology Department, The University of Texas at Tyler; The Department of Pharmaceutical Sciences, Fisch College of Pharmacy, The University of Texas at Tyler

## Abstract

The pathogen *Ophidiomyces ophidiicola* (*O*.*o*.), widely known as the primary cause of snake fungal disease (SFD) has been detected in Texas’s naïve snakes. Our team set out to begin to characterize *O. ophidiicola’s* spread in east Texas. From July 2019 until October 2021, we sampled 176 snakes across east Texas and detected 27 positives cases (qPCR confirmed 27/176). From a ribbon snake with clear clinical display, we isolated and cultured what we believe to be the Texas isolate of *O. ophidiicola*. With over 1/10 snakes that may be infected in East Texas, gives credence to the onset of SFD in Texas.

## Introduction

Across the globe fungal diseases are having large adverse effects on organisms. Fungal pathogens such as *Batrachochytrium dendrobatidis* (BD or Chytrid Fungus) in amphibians and *Pseudogymnoascus destructans* (pathogen in White Nose Syndrome) in bats have been shown to devastate large populations of organisms (Blehert et al. 2009). *Ophidiomyces ophidiicola* (*O. ophidiicola*) has been problematic with specific snake populations, including Eastern Massasaugas Rattlesnakes and Timber Rattlesnakes losing significant portions of their numbers (Allender et al. 2011; Fisher et al. 2012). *Ophidiomyces ophidiicola* has been confirmed as the primary pathogen in Snake Fungal Disease (SFD) (Rajeev et al. 2009). After the initial attachment, *O. ophidiicola* can spread to subdermal tissues causing the host discomfort. Furthermore, individual snakes that do not display clinical signs of the disease can have its normal skin microbiota altered (Allender et al., 2018). Subdermal infection is characterized by lesions on the head, nose, body, and/or tail caused by dysecdysis and may cause face swelling, clouding of the eye, as well as, fluid-filled vesicles (granulomas) (Lorch et al. 2015).

Other effects from SFD include increased basking, and decreased appetite. These are congruent with common symptoms among snakes fighting a variety of infections. Increased basking helps decrease the time between shedding events. These shedding events are a way that reptiles use their innate immune defense to get rid of skin infections (Stromsland et al., 2017). While these processes may help snakes to recover, these modifications make snakes easier targets for predation and more likely to die of malnutrition (Lorch et al., 2015). To date there have not been any known cases of SFD that are verifiably associated with the primary pathogen (*O. ophidiicola*) found in Texas. However, there have been anecdotal sightings of symptomatic individuals in East Texas and fungal infections found associated with *Fusarium* fungus in Southern Texas (Barber et al., 2016). The nearest researched case of SFD was in eastern Louisiana making it a priority to monitor the pathogen’s spread westward (Glorioso et al., 2016). With the addition of climate change increasing the temperature of the planet, these “seasons” will become longer. Increases in temperature will extend the viability of the fungi and geographic range (McKenzie et al., 2018). The purpose of this study is to survey snakes across east and into parts of central Texas to determine if there is a presence of *O. ophidiicola*. This data can aid in determining its prevalence and the populations at risk. This study will track the spread of this pathogen. We hypothesized; that there is a presence of snake fungal disease in Texas. If we find an *O. ophidiicola* isolate from Texas, then we can begin to determine similarities and differences with other strains.

## Materials and Methods

### Encountering Snakes

From May 2020 to November 2021, various methods including shade traps, minnow traps, road cruising, walking and through contact with public (i.e. found snake on their property), were used to create encounters with snakes. Research was conducted in multiple counties across East and Central Texas. We collected samples in Smith, Freestone, Cherokee, Hunt, Navarro, Bell, Washington, Harris, Madison, and Henderson counties in Texas. We also sampled in the Neches River Federal Wildlife Refuge and the Old Sabine Bottom Wildlife Management areas. Shade traps were set out at all locations, however we reserved minnow traps to closer properties. Doing this we heavily reduced the chance of mortality caused by exposure or drowning. We also processed any snakes found on road dead (F.O.R.D.), or sheds we encountered.

Once a snake was encountered, we took down its GPS location, time, date, cover type, the position of the snake and noted any lesions. Additionally, we would use UV flashlights in low light to examine for fluorescence. Vivitrio et al. found in 2021 that the fungal infection often gave off a UV fluorescence. We aimed at looking at that as an indicator as well. Lesions were noted if they were comparable to snakes experimentally challenged in a study by Lorch et al. in 2015. These lesions are defined as thin scales with discoloration, ulcerations, and impacted scales (dysecdysis). Each individual snake was swabbed on the head and then on the body. We would run the swab over the head of the snake five or more times and down their ventral and lateral side five times or more. We did this using one swab for the head and one for the body (Veilleux et al. 2020). Any lesions that were found on the snake were photographed and were individually swabbed. The swabs were stored in a 1.5- or 2-mL centrifuge tube at -20°C until DNA extraction. Any snakes that were handled without gloves on were removed from the data set (n=3, all *O*.*o*. negative).

Nitrile gloves were worn to avoid inadvertent spread of pathogens during handling of snakes. Lysol wipes and 10% bleach solution were used to disinfect equipment and boot bottoms between snakes and sites. Additionally, cleaned and washed clothing was worn on each sampling trip as another safeguard to prevent the spread of the pathogen. The snakes were not clipped or implanted with marking devices. This was done to protect the snakes from the pathogen being introduced directly to their skin. The primary objective of this research is to determine if the disease has spread to Texas and any positive samples are important, regardless of recapture.

### Verification via Quantitative PCR

DNA extraction of the swabs were initiated by soaking them in 1mL of lyticase (300U/mL) in microcentrifuge tubes for 24 hours to help break down the chitin walls of any fungi on the swabs (Bohuski et al. 2015). After the lyticase soak they were put through two freeze/thaw cycles and then centrifuged at 20,000 rpm for an hour to ensure all materials possible were removed from the swabs into the lyticase solution. The swabs were then removed from the remaining solution carefully and placed in another clean microcentrifuge tube and stored in a -80°C freezer for storage. We then put the solution through an extraction (Qiagen DNeasy Blood/Tissue Extraction Kit, Qiagen #69504) using the instructions provided by the kit. The purity and quantity of DNA extracts were determined by nanodrop spectrometry and aliquots of extracted samples were stored at -80°C for further analysis. We used a CFX Connect Real-Time PCR Machine (Bio-Rad, USA) to complete the quantitative polymerase chain reactions (qPCR) to amplify the fungal DNA. The primers used targeted the internal transcribed region (ITS) of ribosomal polymorphisms (Bohuski et al. 2015). We optimized our protocol using negative template and positive control specimens purchased from The UAMH Centre For Global Microfungal Biodiversity (#10769 UAMH). We used the qPCR settings from the methods found in research by Allender et al. 2015 using SYBR Green.

### Ultraviolet fluorescence Measurements and Microscopy

We examined snakes with a UV (365nm) lamp to detect infected areas. Equal amounts of *O. ophidiicola FL* and *O. ophidiicola TX* were loaded into wells of 1% agarose gel and visualized with a standard UV transilluminator to measure the UV fluorescence of the isolates. Due to overlapping emissions, nuclear orange was used to visualize the nuclei of fungi. Nuclear orange LCS1 (AAT Bioquest, #17541) is a non-fluorescent DNA-selective and cell-permeant dye (ex 514nm, em 556nm). Nuclear orange has its fluorescence significantly enhanced upon binding to DNA, reducing background. *O. ophidiicola* was grown in poly-lysine coated 2-well chamber slides (Lab-Tek II, #154461) and then labeled with nuclear orange at 10 μM for 30 minutes. The media was removed and gently washed three times with filtered PBS. After reducing moisture levels, slipcovers mounting was performed with ProLong Gold Antifade reagent. We prepared unstained fungi in the same way minus label to visualize natural UV fluorescence. Samples were analyzed using a Zeiss LSM 880 with Airyscan 5 or an Echo Revolve microscope.

## Results

A total of 176 snakes were encountered during the project spanning 16 species, and a total 27 were positive for the fungal pathogen *O. ophidiicola* (Figure1, Table 1). Using the presence/absence model from Hileman et al. 2018, we’ve categorized these snakes into three groups: SFD Positive (clinical and O.o. positive), Early Stage/Exposed (no clinical signs but O.o. positive), or No SFD/Failed to detect (anything with or without clinical signs that lacked presence of O.o.) (Table 2). Three snakes were removed from the data set due to improper handling.

**Table 1.** Encountered species. Table showing numbers of encountered and qPCR positive snakes by species. *Unknowns are snakes that were deceased and unable to tell the species due to degradation

| SPECIES | NUMBER ENCOUNTERED | O.O. POSITIVE |
| --- | --- | --- |
| *ELAPHE OBSOLETUS* | 35 | 2 |
| *AGKISTRODON PISCIVOROUS* | 20 | 4 |
| *NERODIA ERYTHROGASTER* | 20 | 4 |
| *SHED* | 17 | 0 |
| *THAMNOPHIS PROXIMUS* | 15 | 4 |
| *NERODIA RHOMBIFER* | 14 | 1 |
| *STORERIA DEKAYI* | 14 | 2 |
| *AGKISTRODON CONTORTRIX* | 13 | 2 |
| *MASTICOPHIS FLAGELLUM* | 6 | 2 |
| *HALDEA STRIATULA* | 5 | 2 |
| *NERODIA FASCIATA* | 5 | 1 |
| *FARANCIA ABACURA* | 2 | 0 |
| *LAMPROPELTIS CALLIGASTER* | 2 | 0 |
| *OPHEODRYS AESTIVUS* | 2 | 0 |
| UNKNOWN* | 2 | 0 |
| *LAMPROPELTIS HOLBROOKI* | 1 | 0 |
| *MICRURUS TENER* | 1 | 0 |
| *THAMNOPHIS SIRTALIS* | 1 | 0 |
| *VIRGINIA VALERIAE* | 1 | 0 |
| **TOTAL** | **176** | **27** |

**Table 2.**
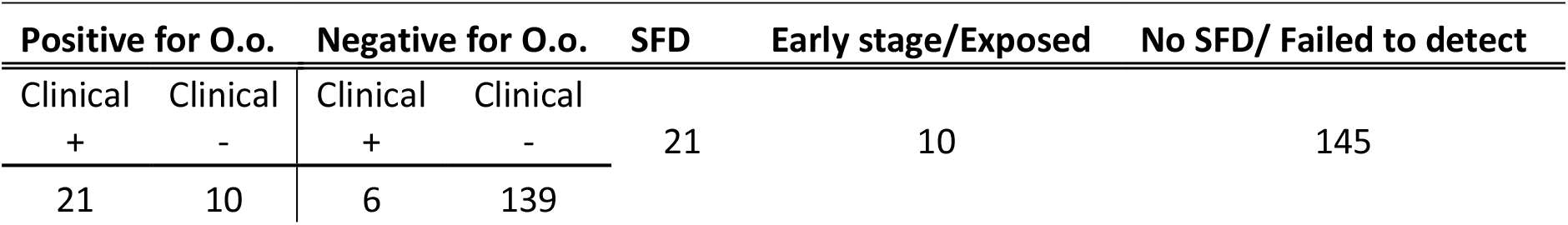
Severity of infection. Table showing infections delineated by metrics created by Hillman et al. 2018. Using clinical and qPCR knowledge to sort snakes into three main categories of SFD, Early stage or Exposed, and No SFD or Failed to detect.

| Positive for O.o. |  | Negative for O.o. |  | SFD | Early stage/Exposed | No SFD/ Failed to detect |
| --- | --- | --- | --- | --- | --- | --- |
| Clinical | Clinical | Clinical | Clinical |  |  |  |
| + | - | + | - | 21 | 10 | 145 |
| 21 | 10 | 6 | 139 |  |  |  |

**Figure 1.**
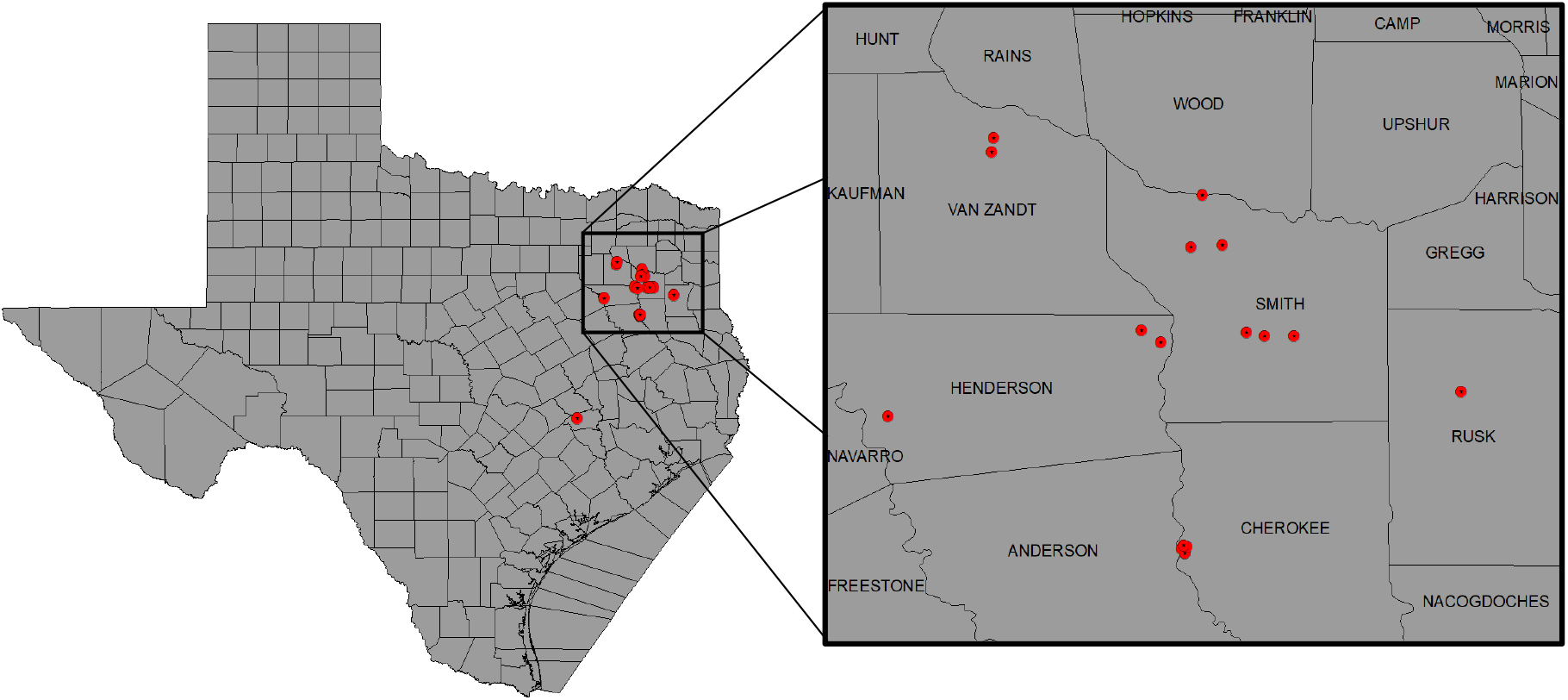
Distribution of *O. ophidiicola* in Texas. Texas State and locations positive for O. ophidiicola are shown as red dots on the map (Table 1).

During our encounters, we found a Western Ribbonsnake (*Thamnophis proximus*) that was in a condition to legally collect and have processed for histology. The snake was found at the OSBWMA, it had mild to moderate crust on its swollen right eye and chin, multiple areas of crusted/missing scales along the ventral side of its body, and dysecdysis on its dorsal side (Figure 2). The snake was moribund and was housed according to IACUC requirements. However, it died approximately two days after collection. Prior to death, we were able to culture fungi from its sores and confirmed via qPCR that it was *O. ophidiicola*.

**Figure 2.**
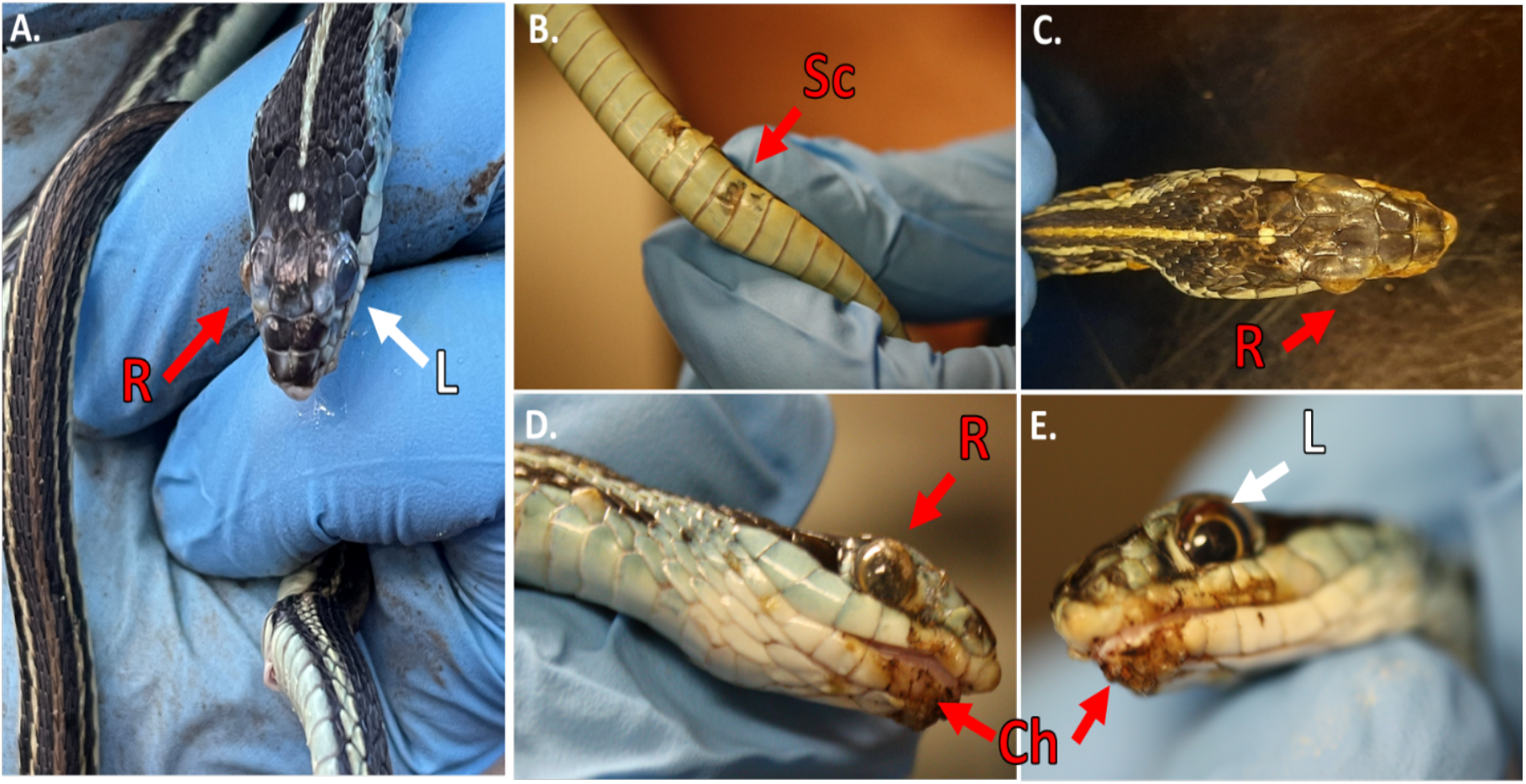
Clinical Signs of Snake Fungal Disease in the Field. Photos of collected Western Ribbonsnake (Thamnophis proximus) A.) Comparison between the gross lesions on the right eye (“R” in red), and unaffected eye (“L” in white) B.) Lesions on ventral scales of the specimen (“Sc” in red) C.) Over the top view of lesions on head D.) Side view of right eye lesions E.) View of gross lesions under the chin (“Ch” in red) of the specimen.

The snake was swabbed again, small clippings taken at the sores, and then the snake was photographed under UV light (Figure3). Afterwards the samples were stored at -20°C until the specimen was delivered to the USGS for analysis. This snake was sent to the U.S. Geological Survey - National Wildlife Health Center (Madison, WI) for processing where Ophidiomyces ophidiicola was isolated in culture from skin lesions and SFD was confirmed histopathologically (Supplemental Material A). Additionally, analysis found that the snake had also been infected by tapeworms. We do not know if this happened before or after infection of the snake by the pathogen.

Long-wave ultraviolet (UV, 365nm) light can detect fungal skin infections and is utilized as a field detection screen tool (Turner et al., 2014; Breyer et al., 2021; Vivirito et al., 2021).,Upon UV exposure, *O. ophidiicola* positive snakes emit a UV fluorescence, as shown with the infected Western ribbon snake from Figure 2 (Figure 3 panels A&B). The intensity of the signal correlates with the amount of fungus at each site (Turner et. al, 2014; Vivirito et al., 2021). To further examine this emission by loading *O. ophidiicola* cultures into the wells of unstained agarose gel. Both *O. ophidiicola TX* and *O. ophidiicola FL* (Figure 3, panel C, lanes 2&3) transmit a detectable signal compared to buffer alone (lane 1) when placed on a standard UV transilluminator (Sung-Jan et al., 2009; Breyer et al., 2021). Furthermore, this ultraviolet fluorescence is detected at the cellular level, as shown with confocal imaging of natural transmission as seen in other fungi including oceanic fungi (Sung-Jan et al., 2009; Breyer et al., 2021) (Figure 3, panel E). UV fluorescence may be the result of high content of chitin and/or ergosterol in fungal species (Turner et. al, 2014; Breyer et al., 2021). Initial swabs from our infected ribbon snake were placed in Sabouraud’s Dextrose medium (SDB) and incubated to test for growth. An isolate from the infected right eye (Figure 4, panel D) was plated on Sabouraud’s Dextrose Agar (SDA) plates and grown under gentamicin and chloramphenicol selection (50 μg/ml each).

**Figure 3.**
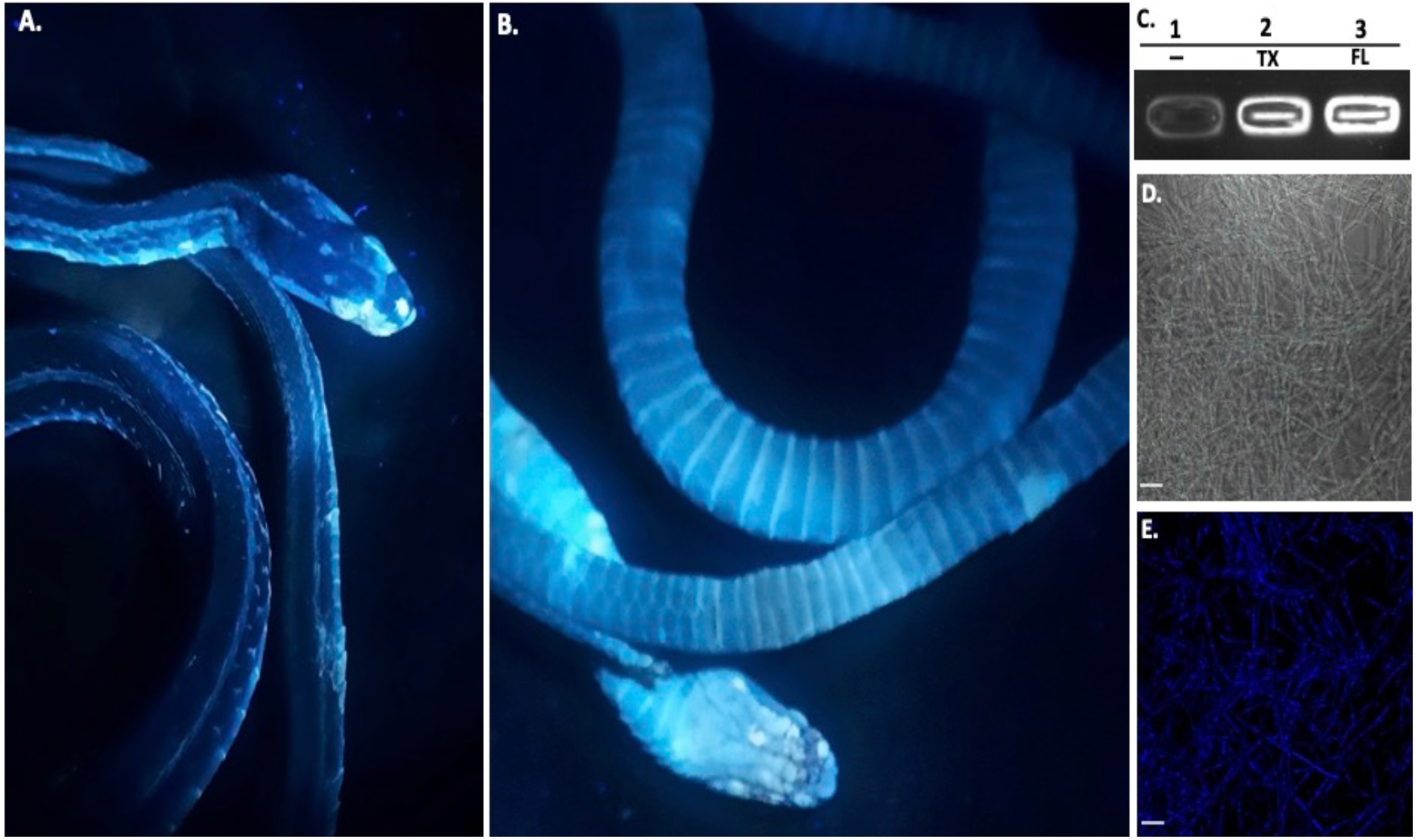
Ultraviolet Fluorescence. Infected Western Ribbonsnake from figure 2 under UV light (365nm) lamp, images shown are dorsal side (A) and ventral side (B). Areas of intensity signal are evidence of the pathogen, *O. ophidiicola*. C.) Transilluminator readings of UV fluorescence between strains *O. ophidiicola FL* (UAMH #10769) and new Isolate *O. ophidiicola TX*. Lane 1. Buffer alone, lane 2 *O. ophidiicola* TX., Lane 3 *O. ophidiicola* FL. (D.) Confocal microscope imagery of *O. ophidiicola* TX in light phase and Texas found strain E.) UV fluorescence after excitement at 358 nm and its emission at 460 nm (blue). Bars: D and E = 30 μm.

**Figure 4.**
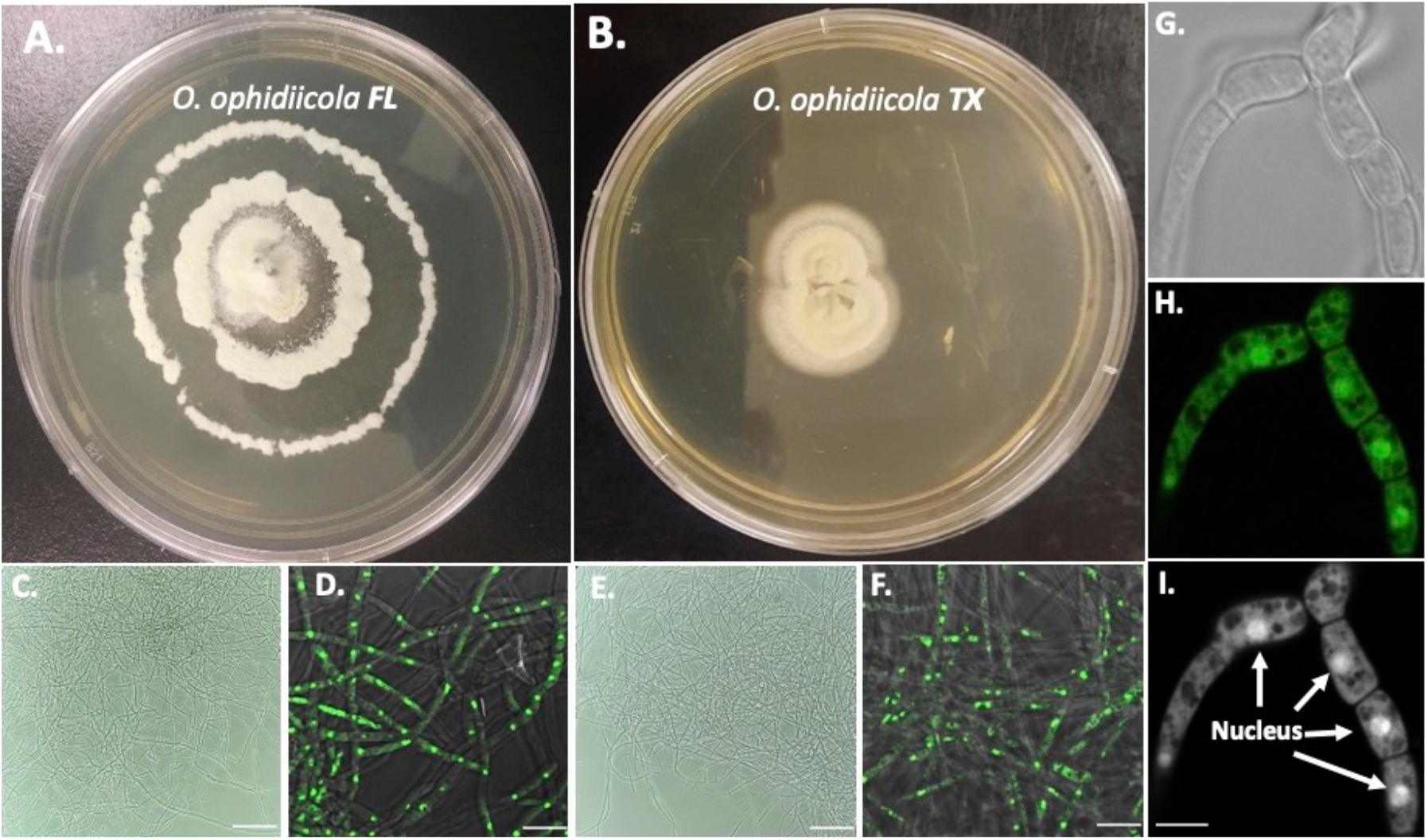
Colony and Microscopic Features of *O. ophidiicola FL* and new Isolate *O. ophidiicola TX*. Comparison of colony formation SDA plates: ***O. ophidiicola FL* (A.)** and ***O. ophidiicola TX* (B.)**. Light microscope images of hyphae phase: *O. ophidiicola FL* (A.) and, *O. ophidiicola TX* (B.). **Nuclear Staining of O. ophidiicola Hyphae**. Comparison of fluorescence between *O. ophidiicola FL* (D) and *O. ophidiicola TX* (F) isolates. Stained with nuclear orange to see the nuclei in the first column and merged with light imagery in the second column. (F) Phase contrast of hyphae branch point. (H) Nuclear orange staining Nuclei in green. (I) Black and white of nuclear staining and defined septate hyphae. Bars: C and E = 150 μm; D and F = 10 μm; I = 5 μm.

Figure 4 shows the comparison growth of *O. ophidiicola FL* (Figure 4, A) and our new isolate, *O. ophidiicola TX* (Figure 4, B). Plates were inoculated in the center and allowed to grow for 25 days. *O. ophidiicola FL* forms a unique bullseyes growth pattern, while *O. ophidiicola TX* stayed in the center with ring halos forming its growth strata (Figure 4, A&B). Examination under a light microscope reveals both are forming a uniform mycelium (Figure 4, C&E). Nuclear orange staining reveals the nucleus within each cell without UV fluorescence interference (Figure 4, D&F). Individual cells within the septated hyphae branches contain a distinct nucleus separation from neighboring cells (Figure 4, G-I). Overall, *O. ophidiicola FL* and our new isolate, *O. ophidiicola TX*, have the same nuclei pattern throughout their hyphae.

## Discussion

The collected snake we found has given evidence to our hypothesis that Snake Fungal Disease caused by *O. ophidiicola* has made it to Texas and has been confirmed using three different avenues. After finding the pathogen in the same area in two consecutive years, on the same species, it is recommended there be a monitoring program for these populations of Ribbonsnakes. Studying these populations in East Texas may give valuable data for documenting spread, and what environmental factors may aid the distribution.

Using UV to aid in detection will be a helpful tool; with more data, we will be able to make these correlations easier and sooner. The ultraviolet light aspect of this fungus adds to the ability to monitor pathogens and are necessary to establish control/mitigation strategies. UV fluorescence is present in other fungi at the cell level responsible for skin infections and forms of marine fungi. The origin of this fungal autofluorescence may result from a high content of chitin or ergosterol in fungal species (Turner et. al, 2014).

This study focused solely on wild snake populations, but one of the primary means of invasive species introduction is the pet trade (Thompson et al. 2018); poor conditions and cramped enclosures are an incubator for pathogens, especially when the animals are a newer monoculture. By monitoring positive cases both in captivity and in the wild we can gain a better understanding of how anthropomorphic and natural based infections spread. Doing so will allow researchers time to come up with solutions to combat this deadly fungus and other pathogens like it.

Our success in retrieving an *O. ophidiicola TX* isolate will allow us to further characterize the cell structures and ultra-structure and then compare them to other isolates. Through nuclear orange staining, we show that each cell has a distinct nucleus and compartments (Figure 4). Their nuclear morphology is a simple circular pattern centered midway in each cell, of whom we estimate the nuclear diameter at between 1.3 and 1.5 nm (Figure 4, G-I). This data begins to characterize the septated hyphae pattern and branch point arrangement of hyphae (Figure 4, G-I). With *O. ophidiicola TX*, we can now fully describe the fungi found in the surrounding areas of Texas, and with further studies gain insights into its growth pattern and infection rate.

## Conclusions

When determining the pathogen’s spread, there are four considerations: behavioral changes in host species, seasonality of the disease, mortality rates attributed to the pathogen, and spatiotemporal patterns (Charron et al., 2013). We have identified that the spread has made it to Texas, giving us a spatiotemporal update on the natural reach of the disease. In future experiments, we will be using our presence/absence data to investigate potential future avenues of dispersal, primarily through the scope of climate change as the area gets more fungal friendly (warmer and wetter). Our team also investigated the UV fluorescence and morphology of the pathogen *O. ophidiicola*, which gave us valuable insights into identification in the field. Increasing accuracy and efficiency of identification will give researchers better tools to find and examine positive snakes. Behavioral changes caused by snake fungal disease have been well documented and were observed during our research. We have yet to determine a seasonality to its onset, but others have shown the condition to be more prevalent in early spring and late summer (McKenzie et al., 2019). These factors are all paramount when tracking the spread of snake fungal disease, and using these fields to bench methods, the more information we can gain in these areas, the more effective we will become at fighting the disease.

## Acknowledgements

Appreciation and gratitude are due to Texas Ecological Laboratories program for funding on this project. Federal Fish and Wildlife along with Leo Gustafson for access to the Neches River Federal Wildlife Refuge. We thank Dr. Jeffrey Lorch, Dr. Jaimie Miller, and other staff at the U.S. Geological Survey - National Wildlife Health Center for assistance with confirming a diagnosis of SFD in a Western Ribbonsnake. Imaging processing was performed by Devon Wade. Texas Parks and Wildlife for access to their properties as well as Dr. Paul Crump. Without the access, funding, and guidance, we could not have finished this experiment.

