## Supplemental Material A for "Snake Fungal Disease Caused by the Fungal Pathogen *Ophidiomyces ophidiicola* in Texas"

### NATIONAL WILDLIFE HEALTH CENTER

**6006 Schroeder Road**
**Madison, Wisconsin 53711-6223**


#### DIAGNOSTIC SERVICES CASE REPORT

**Case:** 30523
**Epi/WID #** 201674

**Supplemental Report** 10/29/2021

**Legal** ☐ **Declassified** ☐ **INV#:**

**Submitter:**
Paul Crump
Texas Parks & Wildlife/Austin
4200 Smith School Road
Austin, TX 78744

**Date Submitted:** 9/9/2021

**Specimen description/Identification/Location:**

| AC | SPECIES | SPECIMEN TYPE | BAND NUMBER | SUBMITTER's ID | COUNTY | STATE |
|---|---|---|---|---|---|---|
| 001 | Snake, Western Ribbon | CARCASS | | | Unknown | TX |

**Diagnosis:**

1) Snake Fungal Disease

**Event History:**

One Western ribbon snake with gross clinical signs of SFD was collected by the University of Texas Biology Department. Samples were PCR positive for Ophidiomyces ophidiicola. Collection date and location not provided.

TPWD would like to submit the snake to verify SFD histologically and culture the fungus.

Legal ☐ Declassified ☐ INV#:

Comment:

**Final Report 10/22/21**

**COMMENTS:**

One adult western ribbon snake of undetermined sex was received for snake fungal disease confirmation. Only an external examination, culture of skin lesions, and histopathology of skin lesions were performed. There was yellow-brown discoloration and mild thickening of the scales around the mouth. There were moderate numbers of yellow-brown discolorations to the ventral scales on the cranial third of the body measuring ~1 x 2 mm. Histopathology shows crusts on sloughing scales on the body and head with suspect fungal hyphae. There is minimal inflammatory response in the sections. Also, there are multiple sections of Plerocercoid cestodes in the subcutis or skeletal muscle with no tissue reaction. Special stains to highlight the fungal hyphae are pending and will be sent in a supplemental report. Fungal culture of the skin yielded *Ophidiomyces ophiodiicola,* which was identified by ITS sequencing. The gross and microscopic findings and culture results are consistent with snake fungal disease (SFD).

Wildlife and Domestic Animal Significance: Snake Fungal Disease (SFD) is a newly-described deep fungal infection of snakes. Investigations are underway to determine whether the fungus *Ophidiomyces ophiodiicola* is the sole cause of SFD or whether it is caused by multiple pathogens. Examinations of snakes with SFD indicate that the fungus invades deeply into the epidermis and dermis, hence molting may not rid the animal of infection despite temporary resolution of clinical signs. The significance of SFD in free-ranging snakes is also currently under investigation. As the name of the disease implies, SFD is only known to affect snakes.

Human Health Considerations: None

Disease Control and Biosecurity: Wear clean disposable gloves when handling sick or dead snakes. Clean supplies and field equipment with soap and water followed by disinfection with a 10% bleach solution (9 parts water, 1 part bleach) between animals and sites. When SFD is already known to occur in a region, snakes whose skin lesions appear to resolve with supportive care and/or antifungal therapy may be candidates for release at their capture site, but these individuals should not be released in an area where the disease has not been previously as it is not known if treated snakes may still harbor viable fungus.

Please continue to monitor this mortality event and provide periodic updates to NWHC Epidemiology Team throughout the course of the event as additional considerations and further investigation may be warranted.

**GROSS, MICROSCOPIC AND DIAGNOSTIC FINDINGS:**

**ACCESSION 001**

**GROSS FINDINGS:**

*External examination*: A thawed 20.41 g adult undetermined sex western ribbon snake is received for snake fungal disease confirmation. The total length is 73.2 cm and the snout to vent length is 53.0 cm. There is yellow-brown discoloration and mild thickening of the scales around the mouth. There are moderate numbers of yellow-brown discolorations to the ventral scales on the cranial third of the body measuring ~1 x 2 mm.

*Internal examination*: Not performed

**MICROSCOPIC FINDINGS:**

1) Skin: Moderate multifocal dermal crusting with intralesional fungal hyphae
2) Skin and head: Marked subcuticual and intramuscular cestodiasis
3) Skin: Suspect single microfilaria
4) Skin: Suspect single parasitic egg (possibly trematode)

**DIAGNOSTIC TEST RESULTS:**

*Microbiology*: Skin, fungal culture: *Ophidiomyces ophiodiicola* was identified by ITS sequencing

**Supplemental Report 10/29/21**

Special staining with GMS for fungal organisms show many fungal hyphae within the crusts on the sloughing scales. Fungal hyphae are 5 um in diameter with parallel walls, acute angle branching, and septations. This is consistent with *Ophidiomyces ophiodiicola* that was cultured from the skin lesions.

**Supplemental Report** 10/29/2021

**Legal** ☐ **Declassified** ☐ **INV#:**

Jaimie L. Miller

Jaimie L. Miller DVM, Dipl. ACVP
Pathologist Fellow

**The USGS-National Wildlife Health Center conducts wildlife disease investigations with state, federal and tribal partners, and we welcome collaborative dissemination of this information (e.g., publication, press release, technical report, etc.). Please contact the pathologist or wildlife disease epidemiologist assigned to this case to ensure that information is accurately interpreted and appropriately credited.**

**Copies To:**

JOHN SILOVSKY
Texas Parks & Wildlife/Austin, 4200 Smith School Road, Austin, TX 78744

This is a Report for your submission to the National Wildlife Health Center.

For consultation regarding diagnostic findings or laboratory testing and results, please contact the pathologist. Contact information can be found underneath the signature line on this report.

For consultation on the significance of this disease to wildlife populations in your area, assistance with disease control and response, or to report field updates (numbers and species affected, geographical distribution, end date, etc.), please contact an NWHC epidemiologist at or 608-270-2480.
